# Paper-Thin Multilayer Microfluidic Devices with Integrated Valves

**DOI:** 10.1101/2020.12.08.416883

**Authors:** Soohong Kim, Gabriel Dorlhiac, Rodrigo Cotrim Chaves, Mansi Zalavadia, Aaron Streets

## Abstract

Integrated valve microfluidics has an unparalleled capability to automate the rapid delivery of fluids at the nanoliter scale for high-throughput biological experimentation. However, multilayer soft lithography, which is used to fabricate valve-microfluidics, produces devices with a minimum thickness of around five millimeters. This form-factor limitation prevents the use of such devices in experiments with limited sample thickness tolerance such as 4-pi microscopy, stimulated Raman scattering microscopy, and many forms of optical or magnetic tweezer applications. We present a new generation of integrated valve microfluidic devices that are less than 300 μm thick, including the cover-glass substrate, that resolves the thickness limitation. This "thin-chip" was fabricated through a novel soft-lithography technique that produces on-chip micro-valves with the same functionality and reliability of traditional thick valve-microfluidic devices despite the orders of magnitude reduction in thickness. We demonstrated the advantage of using our thin-chip over traditional thick devices to automate fluid control while imaging on a high-resolution inverted microscope. First, we demonstrate that the thin-chip provides improved signal to noise when imaging single cells with two-color stimulated Raman scattering (SRS). We then demonstrated how the thin-chip can be used to simultaneously perform on-chip magnetic manipulation of beads and fluorescent imaging. This study reveals the potential of our thin-chip in high-resolution imaging, sorting, and bead capture-based single-cell multi-omics applications.

## Introduction

Integrated microfluidic circuits with elastomeric valves enable automated and programmable manipulation of fluid at the nano-liter scale^1^. Integrated elastomeric microfluidic valves are three-terminal components in fluidic circuits in which a control pressure on one terminal modulates the fluidic current between the other two terminals. In this way, integrated microfluidic devices are analogous to integrated microprocessors where the pneumatic valves binarize the fluid flow as transistors do for current in electronic circuits. These devices are typically fabricated in polydimethylsiloxane (PDMS) and enable accurate and precise reagent delivery with orders of magnitude reduction in reagent consumption. The large-scale fabrication of micro-valves in a PDMS device is enabled by multi-layer soft lithography and allows for complex fluidic work flows to be integrated into a single microfluidic “chip”^2,3^.

On-chip active flow control and automation is particularly powerful for applications like singlecell sorting^4–7^, chemical and enzymatic reactions^2,3,8,9^, and live-cell imaging^10,11^. Firstly, valve-microfluidic chips are advantageous in single-cell flow sorting over simple flow cell devices by performing more complex sorting operations, such as timed multi-channel sorting and solution exchange that can be re-programmed to adapt to various applications. Secondly, on-chip molecular reactions can be programmed as a series of reagent dispensing, mixing, and washing steps^2,3^. The increase in throughput, reliability, and cost savings from reduced reagent consumption has been demonstrated by automated on-chip genomics^3^ and proteomics^8,9^ sample preparation. Finally, valve-microfluidics offer the precision and rapid fluid exchange capability^2,12^ in a programmable package, which can be readily integrated with optical instrumentation for functional live-cell imaging. For these reasons, integrated microfluidic technology offers solutions for high-throughput biology that are not possible with traditional benchtop techniques.

PDMS microfluidic devices are transparent and can be readily incorporated with optical analysis using an inverted microscope. However, several high-resolution optical measurement technologies that require thin samples are incompatible with standard valve-microfluidic chips because standard multilayer soft lithography produces devices that are on the order of centimeters in thickness. These include optical imaging technologies that require high-NA objectives and condensers to deliver excitation light and collect scattered or fluorescent light such as Stimulated Raman Scattering (SRS) microscopy, 4pi confocal microscopy, optical and magnetic tweezers, and Infrared (IR) spectroscopy for enzyme kinetics. Transition SRS microscopy excites the vibrational modes in molecular bonds by focusing ultrafast lasers through an objective and collecting the forward-scattered light with a condenser on the opposite side of the sample^13^. Similarly, optical tweezers that measure molecular interaction forces trap samples with the objective and monitor its motion with a condenser^14^. To achieve diffraction limited resolution in these methods, both the objective and condenser require high numerical aperture (NA) and short working distances. Magnetic tweezers also have small sample thickness tolerance because the low magnetic susceptibility of a single magnetic particle necessitates magnets to get as close to the sample as possible for high-precision control^15,16^. Additionally, the high IR absorption of PDMS and water contributes to background noise in IR spectroscopy, where the resolution indirectly correlates with the PDMS^17,18^. These high-resolution technologies are also unavoidably low-throughput methods and would greatly benefit from the automated fluid control of valve-microfluidics.

In order to accommodate valve-microfluidics to the aforementioned technologies, thin valve-microfluidic chips that can fit into the tight sample space are required. However, the material properties needed for on-chip micro-valves pose a great challenge to the development of such thin multi-layer device. These properties are 1) flexibility for the valves to expand and shut close the flow channel when pressurized, 2) high tensile strength to withstand stretches during alignment, and 3) sufficient rigidity for the top layer to maintain level to the other layer during alignment and resist the torque of the pin connectors during the chip operation. Materials typically used for multilayer microfluidics, including PDMS^19,20^ and hydrogels^21^, meet these conditions when centimeters thick, while 2) and 3) fail at few microns thin. With these limitations, traditional valve-microfluidic chips have been limited to centimeters in thickness. Other thin valve-microfluidic chips have been previously developed by sandwiching a flexible membrane between patterned glass substrates^22,23^. However, these devices require multi-step glass etching which prohibits the benefits of soft-lithography including throughput and rapid prototyping.

To make valve-microfluidics applicable to space-limited experimental configurations, we developed an ultra-thin integrated valve-microfluidic device using a novel fabrication protocol that overcomes the traditional limitations in making thin valve-microfluidic chips. Our “thin-chip”is made of multiple PDMS layers and is under 300 μm thick, including the cover-glass substrate and can be fabricated with standard soft-lithography equipment and techniques. We validate the thin-chip’s compatibility with all of the major valve-microfluidic functionalities including valving, pumping, and input-output control.

We demonstrated application of the thin-chip with high-resolution SRS microscopy of single cells. Confocal Raman microscopy is a powerful imaging technology that acquires high-resolution, chemically-specific images of biological samples without fluorescent labeling, but suffers long acquisition times due to the low cross-section of spontaneous Raman scattering. SRS greatly enhances the signal for Raman microscopy to enable imaging live samples at video rate^24^. This technique achieves signal enhancement through a nonlinear interaction using two pulsed lasers, with center frequencies chosen such the difference frequency matches a vibrational frequency of interest. SRS provides reduced non-resonant background compared to coherent anti-Stokes Raman scattering microscopy (CARS), however SRS is susceptible to additional background from nonlinear interactions such as cross-phase modulation (XPM), a process where the two lasers affect the sample’s refractive indices for each other^25,26^. XPM leads to unwanted intensity variations across the image, as the additional refraction can cause clipping on optical elements. Such intensity variations obscure the signal of interest and generate non-chemically specific signal at wavenumbers that do not correspond to any molecular vibrations present in the sample. While XPM can be avoided through matching the NA of the condenser with the objective, the short working distances of the high NA lenses limits the sample thickness to under 1.5 mm, thus restricting the use typical valve-microfluidic chips.

In this study, we demonstrate the integration of thin valve-microfluidic chips with high-resolution SRS and we present improvements in the background noise reduction and removal of unwanted signals with the thin-chip.

We also demonstrate the thin-chip’s enhancement of on-chip magnetic bead control on apparatuses that combine an inverted microscope with magnets. On these setups, magnets can only be positioned on top of the sample to allow imaging. Since the magnetic force is inversely proportional to the square of the distance between two magnetic objects, traditional valve-microfluidic chips could not utilize for these applications due to its thickness weakening the magnetic force. With its reduced thickness, the thin-chip improves magnetic bead control inside valve-microfluidic chips by allowing magnetic tweezers to come within 100 microns of the sample.

Improved bead control not only aids magnetic tweezing applications, but can also create packed bead columns to facilitate on-chip molecular selection. The ability to immobilize single functional beads in the presence of fluid flow can also be used to grab and sort single cells. While bead columns can also be created in thick chips by filter valves^3^, they require new device designs, whereas in the thin-chip, beads columns can be created in any existing design by using magnets. Bead based cell sorting is also not possible with filter valves because both the beads and the cells are larger than the filter valve pores that are used for straining. Here, we validated the advantage of our thin valve chips over thick chips in manipulating beads in the device by comparing the ability to immobilize two different sized beads in various conditions.

### Multilayer soft-lithography for thin-chip fabrication

Our goal was to engineer a valve-microfluidic chip that operates between two high NA lenses with a combined working distance of 1.5 mm. After subtracting the 150 to 170 μm thickness of the required cover-glass, we were limited to roughly 1 mm thickness that includes all layers. Additionally, the device needed to accommodate pin connectors for valve control and sample input. These requirements must both be met in a functional thin-chip, however the imaging portion demands millimeter thickness while the pin connector requires centimeters thickness.

To meet these design specifications, we designed a device partitioned into different regions with multiple thicknesses. The thin area called the ‘imaging region’ is under 300 μm in thickness and the two thick areas named ‘bus’ are centimeters thick (Fig 1a and b). With this approach the two different regions can independently satisfy the two opposing requirements for the thin valvemicrofluidic chip.

**Figure 1.**
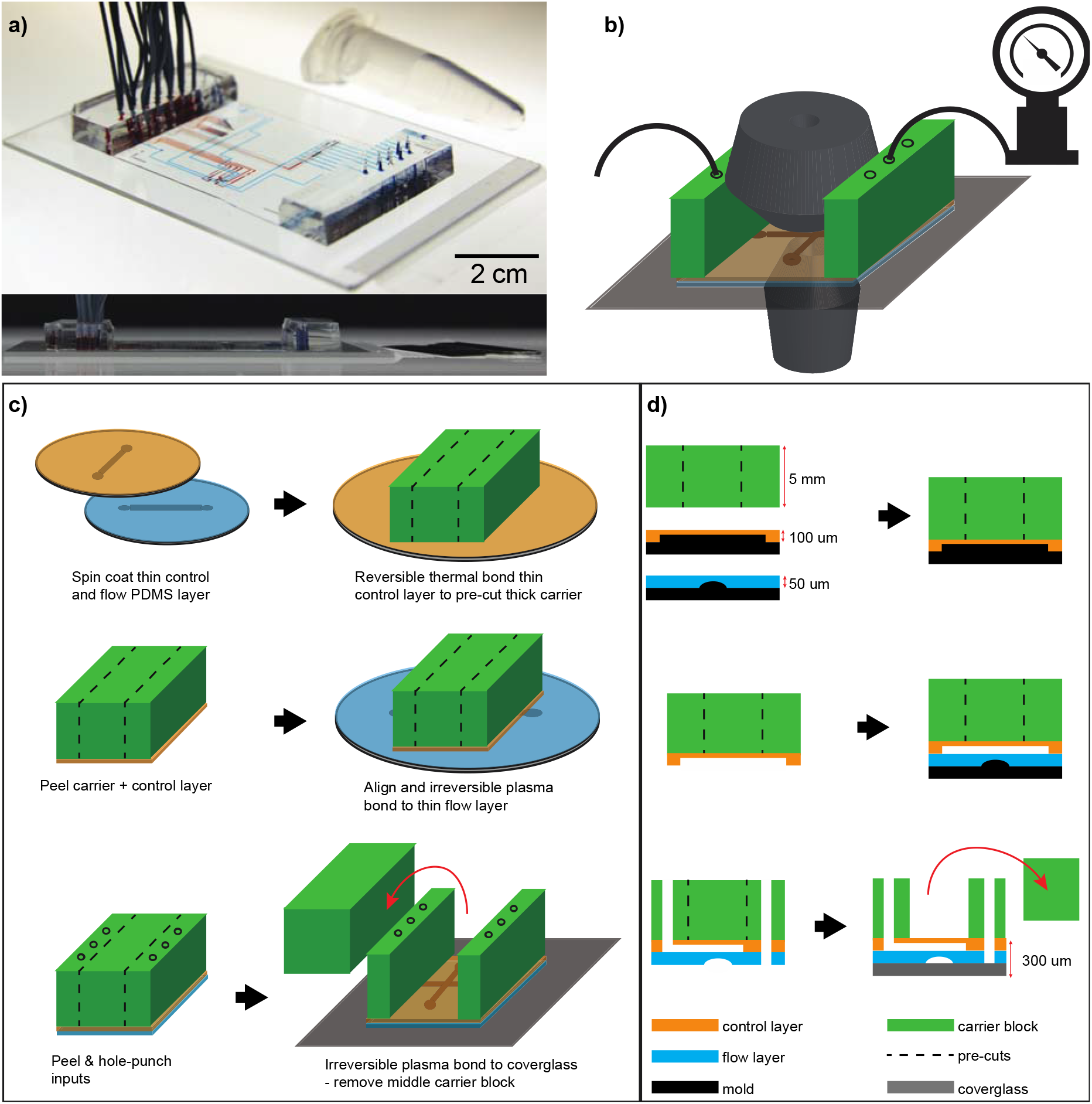
The thin valve-microfluidic chip. **a)** The angled (top) and side (bottom) view of the thin-chip with food coloring in the control channels with a 1.5 mL Eppendorf tube (top) and a razor blade (bottom) for size comparison. The input bus (thick region) is ~1 centimeter and the viewing window (thin region) is < 250 μm thick (2 cm scale bar in photo). **b)** A schematic of the thin-chip loaded between a high NA condenser and objective on an inverted microscope. **c)** A schematic of the thin-chip fabrication protocol in 3D and **d)** in 2D xz cross section. The layers and features are labeled in d).

Our strategy to handle and align two thin layers was to utilize a thick ‘carrier’ layer with a pre-cut detachable block (Fig 1c and d) during the fabrication process. The carrier layer was designed to bond and transfer the thin valve layer to the thin flow layer while providing rigidity and support during alignment. After alignment and final bonding to the glass substrate, a portion of the carrier layer was detached to expose the thin viewing window. The remaining carrier region would become the bus for the pin connector interface (Fig 1a).

Designing this protocol required optimizing the relative bonding strength between the multiple layers so that thin control and flow layers can be effectively transferred from the mold to the substrate, and the carrier layer can be selectively removed. Specifically, peeling the detachment block from the carrier layer requires that only the carrier layer is removed while the two thin layers below remains attached to the glass (Fig 1c and d). To achieve this, we compared various bonding methods and parameters including thermal, plasma, and adhesive bonding and found the most robust solution.

Bond strengths between two different PDMS layers and glass can be tuned by adjusting the bonding method such as plasma or thermal bonding as well as controlling parameters like gas type, gas flow rate, plasma RF power^27^, and baking temperature and duration. Optimized plasma and thermal bonding methods both generate bonds capable of withstanding valve control pressures of above 30 PSI. However, the uniformity and density of covalent bonds created through plasma bonding grants significantly stronger bonds than that of the thermal bonding. Taking advantage of this difference, we plasma bonded the thin layers to each other and to the glass substrate for permanent bonding while thermally bonding the carrier layer to the top thin layer to remove the detachable block. For fabrication reproducibility, we designed our protocol to use only the parameters that maximize bonding strengths for a given bonding method to give tolerance to PDMS batch effects and varying lab conditions

Cover-glass substrates are required for high NA objectives; however, large thin cover-glasses are fragile and difficult to handle. PDMS chips bonded to cover-glass substrates can easily crack from the torque generated by the weight of the connected tubes. The fragile nature of cover-glass posed another serious challenge to fabricating the thin-chip during the detachment region peeling process as the peeling force cracked the glass. As a solution, we created an acrylic adapter and bonded it to our thin device with an adhesive for the needed sturdiness. Both the acrylic and adhesive film were laser cut and aligned to avoid masking of the viewing window (supplementary figure 1). The adapter was also bigger than the cover-glass to support the entirety of the chip.

The thin-chip used in this study was designed to isolate, image, and sort single particles and cells (supplementary figure 2). The device has three sample and buffer inputs, each, and is designed with eight sample outputs to interface with a column of a 96-well plate. To reduce the number of control channels, a flow multiplexer is integrated into the design^2^. The imaging region is designed with an H-bridge fluid circuitry that reverses the flow direction to switch between the sample collection and waste modes. This design also minimizes the cell input and collection buffer channel overlap to prevent cross contamination. Using this chip, we achieved deterministic active sorting, actuated by complex high-resolution SRS imaging. After imaging, the cells were redirected to the sample outlets for collection or the waste channel.

### Thin-chip performance characterization

We compared the valve actuation speed between thin and thick devices with the same design. Valve actuation kinetics were determined by loading fluorescent dye in the device’s flow channels and measuring the cumulative fluorescence in the flow channel directly beneath a valve transitions between open and closed states. The channel was the brightest at the open state and darkest at the closed state since all the dye solution was pushed out of the flow channel when the valve is compressed. The fluorescence was measured by focusing the excitation beam on the center of the flow channel and the fluorescence emission was continuously collected with photomultiplier tube (PMT) with microsecond temporal resolution. We found that the thin-chip actuation rates were comparable to a traditional thick device (Fig 2a).

**Figure 2.**
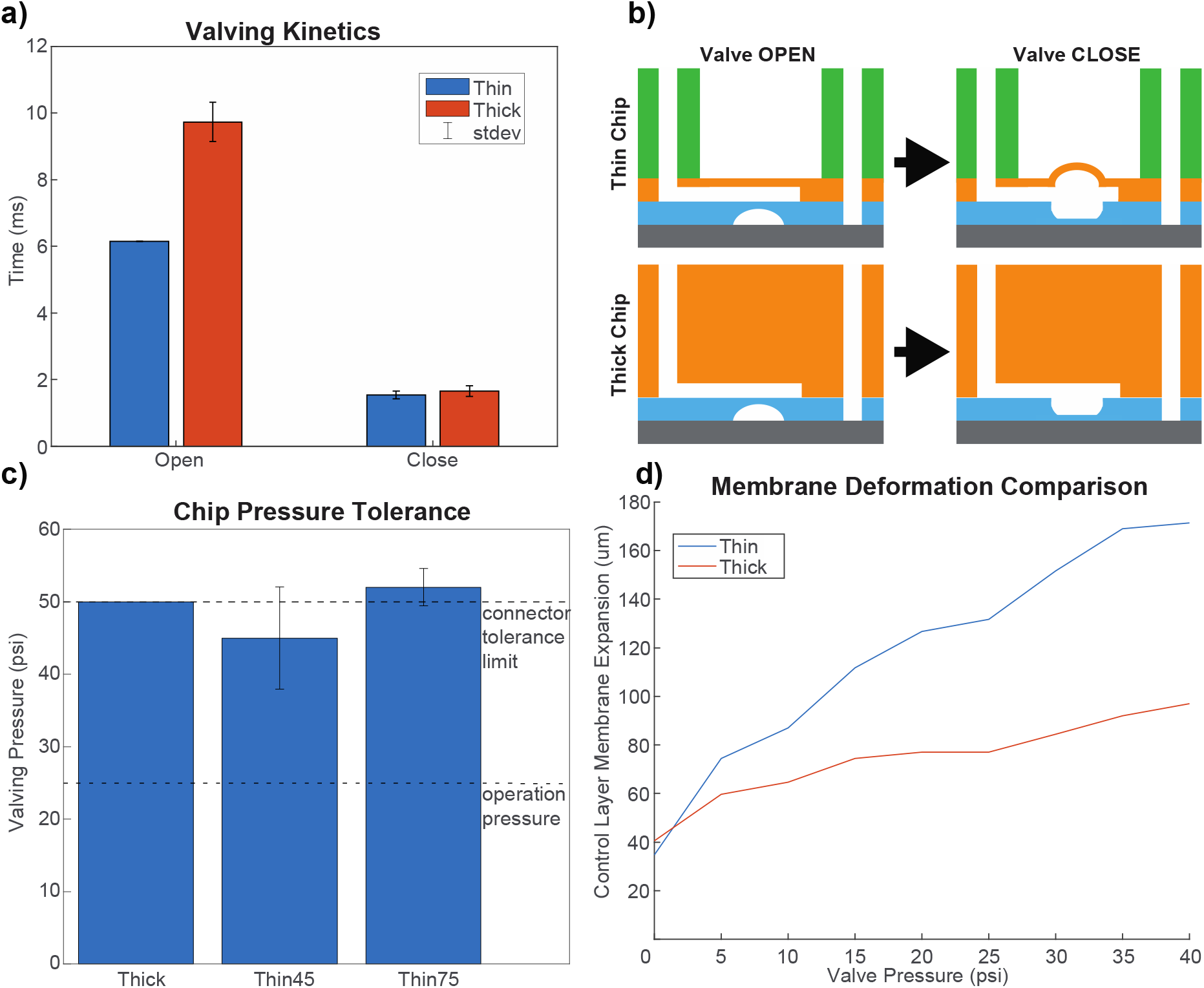
Characterization of the thin-chip in comparison to the traditional thick chip. **a)** Valve opening and closing speed averaged over five operations. Black line denotes standard deviation of the mean error bar. **b)** A cartoon of the 2D xz cross section of the thick and thin-chip during valve operation (the color scheme is as follows: green, orange, and blue are the carrier, control, flow layers, respectively, and grey is the glass substrate. **c)** The chip pressure tolerance averaged over ten chips each with the black line indicating standard deviation of the mean. The bottom dotted line indicates the 25 PSI operation pressure and the top line indicates the pressure tolerance of the seal between the pin connector and input hole. Thin45 and Thin75 denotes thin-chips with 45 μm and 75 μm control layer thicknesses, respectively. **d)** The degree of PDMS membrane expansion comparison between the thin and thick chip when pressurized for valving.

Next, we compared the control pressure tolerance between the thin-chip and traditional thick-chips. Common failure modes of microfluidic chips at high valve pressures are delamination of the PDMS-glass and PDMS-PDMS bonds and connector pin/PDMS unsealing. Additionally, the valves on the thin-chips can ‘balloon’ and rupture upon pressurization due to high PDMS membrane flexibility at such low thickness (Fig 2b). To compare control pressure tolerance, we fabricated multiple thick and thin-chips with 45 μm and 75 μm control layer thicknesses and increased the pressure at 5 PSI increments until the leakage occurred. Both the thick and thin-chips with 75 μm valve layer held above 50 PSI and only failed at the pin/PDMS connection with no delamination, while the thin-chips with 45 μm valves ruptured at 40 PSI (Fig 2c). Thus, there was no substantial difference in pressure tolerance between the thick and 75 μm thin-chips. The total PDMS thickness of the 75 μm thin-chips were 120 μm and under 300 μm including the cover-glass substrate. These chips fit easily between two high NA objectives and the operation pressure of 25 PSI is well below the valving pressure limits.

One major difference between the thin and the thick chips was the amount of membrane deformation when pressurized for valving. This difference is a result of the higher elasticity of the top PDMS layer in the thin-chip in contrast to the thick chip. We quantified the amount of PDMS expansion during valving as a function of varying valve pressures. We measured this by scanning the valve with a focused infrared beam spot in Z and increasing the valving pressure while collecting PDMS SRS (supplementary figure 3). In our comparison, the membrane on top of the valves in the thin-chip expands twice as much as the thick chip (Fig 2d), but this did not affect the valve actuation performance or reproducibility.

### Thin-chip facilitates SRS Imaging

Thin-chips can improve SRS microscopy performance when imaging cells in a microfluidic device because of compatibility with the short-working distance objectives and condensers required in high-resolution SRS microscopy. To demonstrate this improvement in SRS imaging, we imaged mouse N2a cells inside the trapping chamber of a thin-chip designed for single-cell sorting. We compared these images to SRS microscopy in a traditional PDMS device images with a long working distance condenser, by placing a 1 cm PDMS block on top of the thin-chip and re-imaging the cells (supplementary figure 4). In this way we could directly compare images of the same cells between the thin and thick chip configuration. We imaged at two wavenumbers, 2950 cm^−1^, corresponding to CH_3_ stretches, with most of the contributing signal coming from proteins with high fraction CH_3_ bonds, and at 2135 cm^−1^ as a negative control since no mammalian biomolecule correspond to that vibrational frequency. All images were acquired using a 1.2 NA water immersion objective. For images acquired in the thin-chip a 1.4 NA oil immersion condenser was used with a working distance of 1.6 mm. For the thick chip a longer 5.4 mm working distance air condenser was used, with a NA of 0.8. As is apparent in figure 3a, signal exists in the 2135 cm^−1^ channel in the thick chip, reminiscent of a phase-contrast image, which corresponds to the unwanted XPM signal. This signal is absent in the images acquired using the thin-chip and oil condenser. Additionally, the overall background level for the images acquired with the thick chip in the 2950 cm^−1^ channel is higher than for the thin-chip, due to XPM signal contributions from the buffer (Fig 3b). More quantitatively this can be characterized by comparing the information of the images in both channels, for both chips, using a measure of the entropy of the image. Higher entropy, corresponding to higher information content, can be seen at borders of the cells in the 2135 cm^−1^ images for the thick chip as compared to the thin-chip (figure 3c and supplementary figure 5). Additionally, while overall the information content of the 2950 cm^−1^ channels is comparable between the two chips, there are areas of the thin-chip images which reach higher entropy levels, indicating greater intensity variation in these regions.

**Figure 3.**
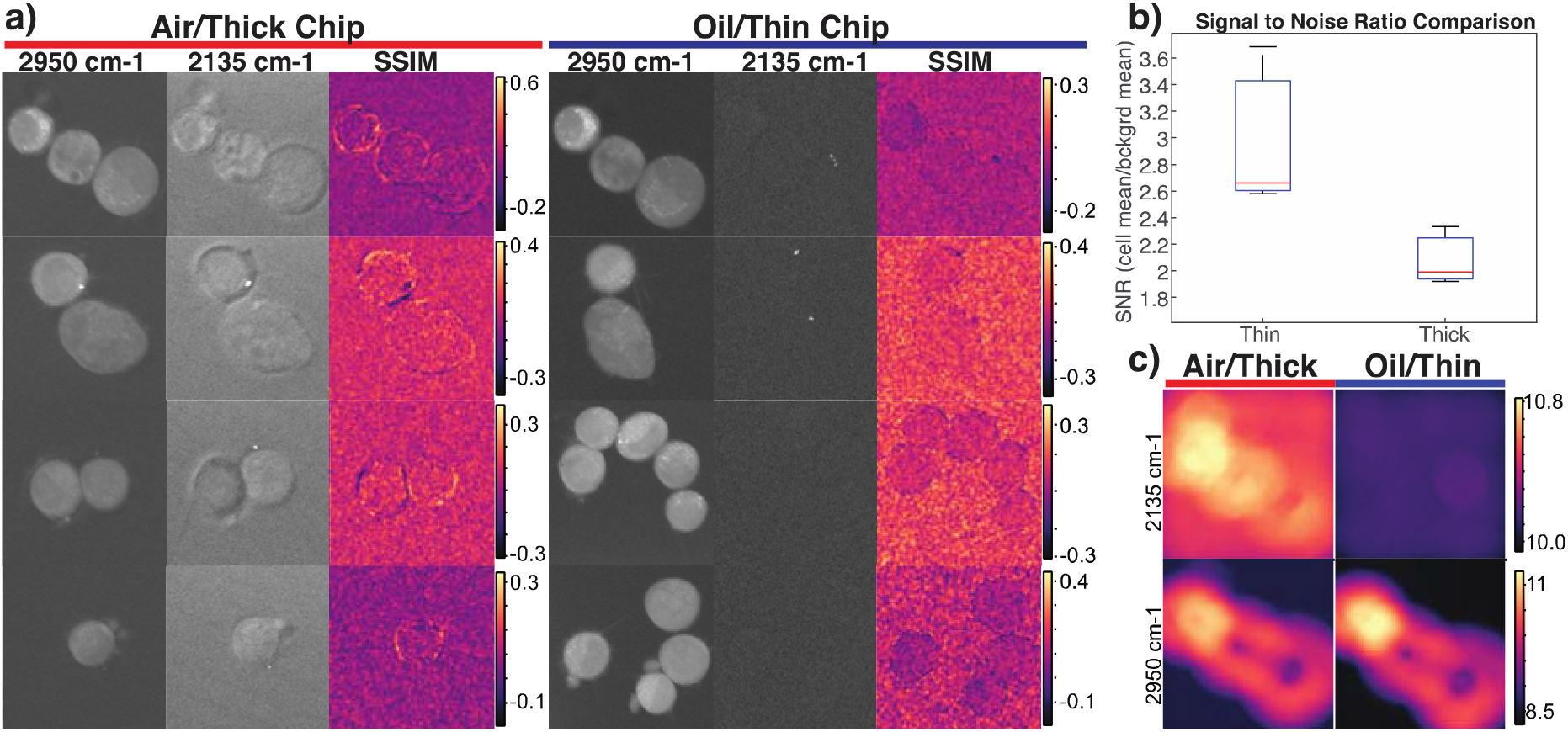
Single cell two color SRS imaging using the thin and thick chip. **a)** SRS images acquired at 2950 cm^−1^ and 2135 cm^−1^ with corresponding local structural similarity index metric (SSIM) images comparing similarity between the two channels. The SSIM metric was calculated pixel by pixel over a local region using 3-pixel standard deviation gaussian weights. The three columns on the right correspond to images acquired using the thick chip and a 0.8 NA air condenser, while those on the left correspond to the thin-chip using a 1.4 NA oil immersion condenser. **b)** The signal to noise ratio (SNR) between images collected of the same cells in the thin and thick chips, calculated as the mean intensity of the cell areas over the mean intensity of the background. **c)** Local entropy maps for the 2135 cm^−1^ (top) and 2950 cm^−1^ (bottom) images, acquired with the thick chip (left) and thin-chip (right) of identical cells. The entropy gives a measure of the number of bits needed to encode the information present in an area surrounding each pixel. An 80-pixel circle was used for the area.

We furthermore characterized the structural similarity between the images acquired at the two wavenumbers, using the structural similarity index measure (SSIM). For images acquired in the absence of the unwanted XPM signal, the structural similarity should be very low between these channels, as the 2135 cm^−1^ channel should contain only noise. However, there is relatively high structural similarity at the edges of the cells in the thick chip using the air condenser, indicating similar feature content (Fig 3a). Note that while it seems that there is cell contrast in the SSIM images for the thin-chip and oil condenser, the metric actually indicates *lower* shared structure due to the transition from just noisy background to internal cell signal.

### Magnetic bead manipulation in in the thin-chip

To verify the potential of sorting single cells with antibody coated magnetic beads using our thin-chip, we validated our ability to immobilize single magnetic beads in a flow field. For this test, we imaged fluorescently labeled 3 μm magnetic beads in the flow channel at varying flow pressure while positioning a rare earth magnet above the trapping chamber (Fig 4a). In our experiment, we successfully immobilized single magnetic beads and resisted up to 2.5 PSI flow pressure by placing the magnet directly on top of the chip. In contrast, the beads were unable to resist even 0.5 PSI when the magnet was lifted to above 1 mm (Fig 4b and supplementary video 1). This result shows the clear advantage of using the thin-chip to control and immobilize single beads in the presence of flow that posed a challenge in traditional valve-microfluidic due to its centimeter thickness and the low magnetic susceptibility of microbeads.

**Figure 4.**
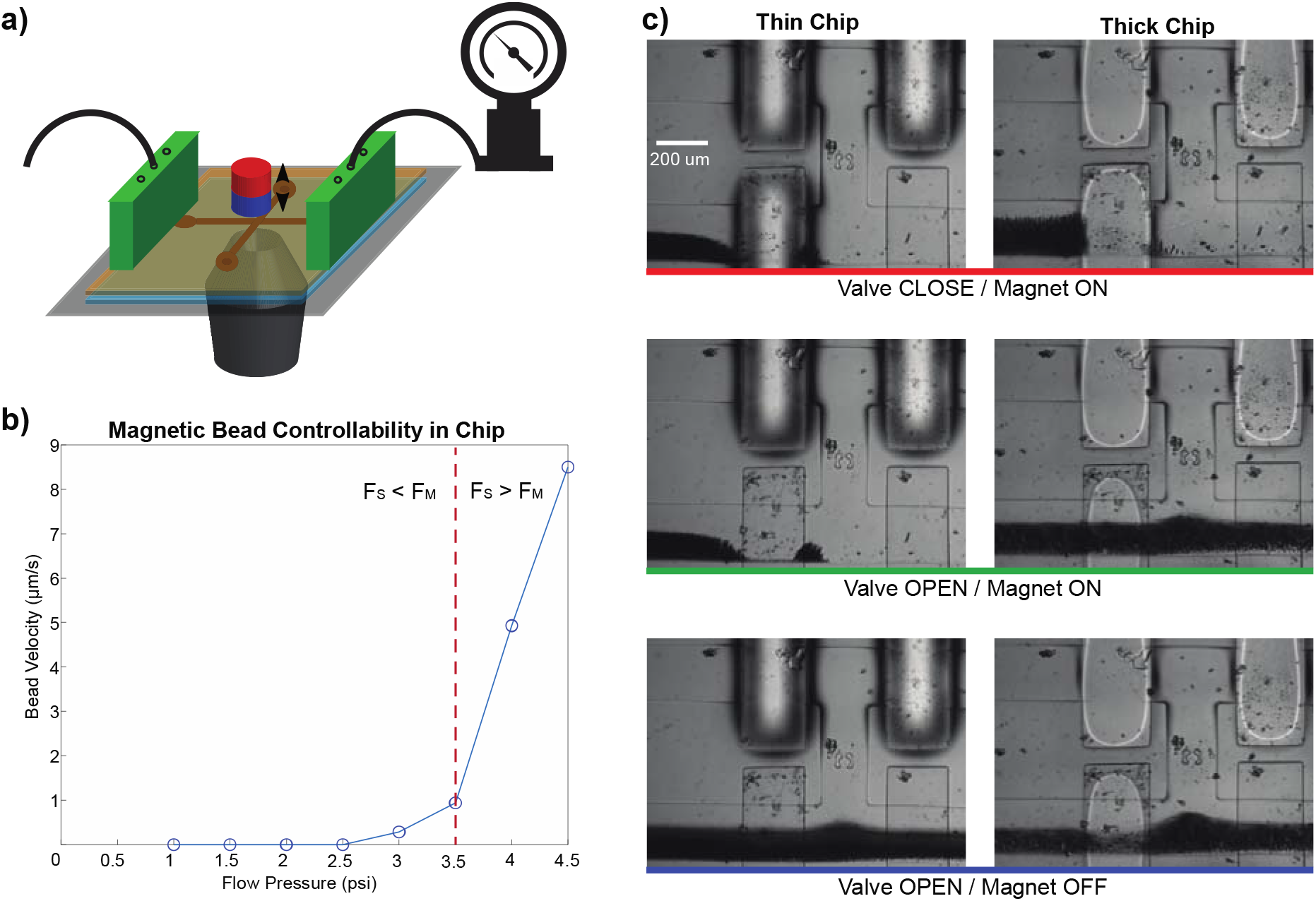
Magnetic bead control demonstration on valve-microfluidic chips. **a)** A cartoon of the experimental setup where the thin-chip was loaded on an inverted microscope. The compact magnet with manual height position control was placed on top of chip. Magnetic beads were injected into the chip by a tube and a regulated external pressure source. **b)** A plot of 3 μm magnetic bead velocity at varying flow pressures in the thin-chip with a stationary magnet directly on top. The red dotted line indicates the threshold where the shear force (F_S_) dominated the magnetic force (F_M_) that held the beads in place and resisted flow. **c)** Reversible 1 μm SPRI bead column creation in the flow channel of the chips with an external magnet. The three images in each column are sequential snap shots of the flow channel with changes in valve states and magnet placement. The external magnet was placed slightly off-center to enable top illumination.

Next, we aimed to demonstrate our ability to create reversible magnetic bead columns in our thin-chip using a common magnet. For this validation, we imaged the flow channel of the thin-chip while flowing 1 μm SPRI beads at 5 to 10 PSI with and without the of the external magnet placed on top of the chip. To compare this with the thick chip, we performed the same experiment with a 1 cm thick PDMS block placed between the magnet and the surface of the thin-chip (supplementary figure 4). Our results demonstrate that the magnetic bead columns can only be generated in the flow channels of the thin-chip while simultaneously imaging with an inverted fluorescent microscope (Fig 4c).

## Discussion

We developed a new generation of paper-thin integrated valve-microfluidic devices with a thickness of less than 300 μm. We demonstrated that the thin-chip valves displayed comparable functionality, reliability and operation speed to traditional multilayer integrated devices with orders of magnitude reduction in thickness. We demonstrated the advantages of our thin-chip with two-color SRS imaging of live single cells, where our thin-chip generated superior image quality with increased resolution and signal-to-noise ratio compared to traditional thick chips. Combining SRS imaging with integrated valve-microfluidics enables many new applications, including single-cell sorting based on high-resolution hyper-spectral SRS imaging. Joined with RNA sequencing, this can add information about chemical composition and morphological information to single cell profiles^28^ for discovering new cell types and related marker genes.

Using our thin-chip, we also demonstrated the ability to simultaneously perform high-magnification imaging while manipulating magnetic beads on an integrated valve-microfluidic chip. The ability to automate and program solution exchange steps will greatly benefit the field of magnetic tweezers. Although magnetic tweezing is a powerful platform to study conformational dynamics and forces that act on single-molecule, these experiments are inherently low-throughput which limits the number of conditions and replicates that can be performed in an experiment. Because biomolecules interact with many different molecules in an extremely large chemical space in physiological conditions, it is impossible to manually quantify each interaction at a meaningful scale^29^. With the thin-chip, the magnetic tweezing community can take advantage of the automated large-scale experimental capability of valve-microfluidics.

The ability to immobilize magnetic beads in the thin-chip while being compatible with inverted microscopes also opens new possibilities for high dimensional multi-omics measurement of single cells. For example, reversible bead columns can be produced for protein or DNA purification by packing magnetic beads in the flow channel with an external magnet. A single antibody-coated bead can also be immobilized in the thin-chip and be used to bind and sort single cells based on their surface proteins, as “programmable” microfluidic pillar arrays^30,31^. However, the immobilized bead-based cell sorting is more translatable because it can be readily implemented and integrated into existing chips. This approach can also detect vastly more cell markers by simply changing the beads during a single chip run. In combination with both imaging and downstream genomic analysis, such a method can enable multidimensional cell profiling.

Finally, the flexibility of the thin-chip technology also enables the possibility of implementing microfluidic valves and active fluidic control in the rapidly growing field of flexible-thin microfluidics for wearable devices. For example, recent PDMS-based sweat sensors could implement integrated valves for multiplexed assays using the thin-chip technology described here^32,33^. Paper-thin valve-microfluidic devices combine the advantages of integrated microfluidic circuits with the thin form-factor of integrated electronic circuits representing a major step in the application of microfluidic large-scale integration.

## Acknowledgements

We thank Annie Maslan, Anushka Gupta, Dr. Nicolas Altemose, and other members of the Streets Lab for helpful discussions. This work was supported in part by National Natural Science Foundation of China (81950410636), and the Li Dak Sum Yip Yio Chin Kenneth Li Marine Biopharmaceutical Development Fund. Aaron Streets is a Chan Zuckerberg Biohub investigator.

## Methods

### Microfabrication

The thin chip was fabricated at UC Berkeley by soft lithography, where PDMS was casted on a photoresist patterned silicon wafer mold. Three separate molds were used to make the 1 cm carrier, 45 μm control, and the 75 μm flow layer. The carrier was pre-cut and partially cured before thermal bonding to the control layer. The control layer on the carrier was then manually aligned and oxygen plasma bonded to the flow layer. After hole punching the input and output ports, the three-layer PDMS chip was then plasma bonded to a number 1.5 cover-glass. Finally, the resulting product was bonded to a laser cut acrylic adapter and the removable middle portion of the carrier was peeled to reveal the 110 μm thick imaging region.

The molds were fabricated using standard micron photolithography. The flow layer mold was patterned in three steps: (1) rounded 30 μm, (2) rectangular 5 μm, and (3) rectangular 30 μm features. The rounded features were produced by spin-coating AZ40XT photoresist (IMM) on a Hexamethyldisilazane (Sigma) coated 4 inch silicon wafer (University Wafers), patterning with 365 nm ultraviolet exposure (OAI 206) through a mask designed (AutoCAD) and printed at 20,000 dpi (CAD/ART), developing in the 300 MIF developer (IMM), then slowly heating from 65°C to 190°C with a 10°C /hour ramp. The rectangular features were serially added to the mold by spin-coating, aligning, and UV exposing SU-8 2005 photoresist (Kayaku) to 5 μm and SU-8 3025 (Kayaku) to 30 μm then developing in SU-8 developer (Kayaku). The control and carrier layers were single layer molds that were fabricated by spin-coating 30 μm of SU-8 3025 photoresist on a silicon wafer, followed by UV exposing and developing. The molds were treated with trichloromethylsilane (Sigma) before soft lithography.

The PDMS chip fabrication was done with standard on-ratio bonding multilayer soft lithography protocols. PDMS (Momentive) was mixed at a 10:1 silicone to crosslinker ratio and degassed on an orbital mixer (Thinky AR250), poured onto the carrier mold, and partially cured by baking at 70°C for 24 minutes. The carrier was pre-cut to 1 mm from the top by placing two 1 mm substrates (glass slides) next to the top-side-down carrier then cutting with a razor larger than the gap between the two substrates, which leaves an uncut 1 mm thickness. The same PDMS mix was spin-coated on the control and flow layer mold to 45 μm and 75 μm, respectively, and baked at 70°C for 14 minutes. The carrier was manually aligned to the control layer and thermally bonded by baking at 70°C for 5 minutes. The control layer on the carrier was plasma bonded to the flow layer by exposing both to oxygen plasma (brand) for 10 seconds at 0.283 mTorr with X power, then manually aligning under a stereomicroscope (Nikon). After peeling the bonded three PDMS layers from the flow layer mold, the chip was hole punched (Syneo, ID 660 μm tips) on the designated inlet/outlet ports. The chip was then plasma (brand) bonded to a number 1.5 cover-glass (Thomas Scientific) by exposing to oxygen plasma for 22 seconds at 0.283 mTorr with 4% (X) power. The entire assembly was then bonded to a laser cut ½ inch acrylic adapter with a double-sided tape (3M 468MP). The middle portion of the carrier was removed by cutting the 1 mm remaining portion on top with a razor blade then carefully peeling off. The final product was baked at 70°C for 2 hours.

### Microfluidic controller and chip operation

The flow and control pressures were activated by KATARA (http://streetslab.berkeley.edu/tools/katara/) that controlled array of solenoid valves (Pneumadyne) connected to the computer by a USB interface (Arduino Mega).

The valve operation pressure was 25 psi and the flow pressure were varied between 0.5 to 10 PSI using a high-resolution manual pressure regulator. The surface of the flow channels in the microfluidic chip was passivated with 0.5% w/v Pluronic F-127 (Sigma) in water for 10 minutes to reduce bead and cell adsorption.

### Microscopy

The synchronized dual output of a femtosecond oscillator/OPO (Insight DS+, Spectra-Physics) provided the pump (tunable OPO output), and Stokes beams (fundamental at 1040 nm) for the experiment. The power of both beams was controlled using two variable attenuators consisting of a half-wave Fresnel rhomb (for the pump), or half-wave plate (for the Stokes), and a polarizer. The pump beam was tuned to 796 nm for CH_2_/CH_3_ imaging, and 851 nm for off-resonance imaging, corresponding to Raman shifts of 2950 cm^−1^ and 2135 cm^−1^. The actual spectral bandwidth was ~160 cm^−1^ due to the short pulse duration of ~120 fs. The output from the Stokes beam was then intensity modulated at 10.28 MHz using a quarter waveplate (Thorlabs), a resonant electro-optic modulator (EOM; Thorlabs) and a Glan-laser polarizer (Thorlabs). The resonant EOM was driven by a function generator (33120A, Hewlett Packard), and an additional power amplifier (ZHL-32A+, Minicircuits). The pump beam path length was controlled by an optical delay line (FCL200, Newport) to ensure coincident arrival of the pulses of the two beams. The beams were then combined on a 1000 nm short-pass dichroic mirror (Thorlabs), and fed into the scan head of a confocal scanning microscope (FV1200, Olympus). The beams were delivered to the sample by a near infrared optimized 60x water immersion objective with a numerical aperture (NA) of 1.2 (UPLSAPO60XWIR, Olympus). After the sample, the lasers were collected by a 1.4 NA oil condenser (CSC1003, Thorlabs; or equivalently D-CUO, Nikon) or 0.8 NA air condenser. The Stokes beam was then filtered by a 1000 nm shortpass filter (Thorlabs). A series of relay lenses delivered the pump beam to a photodiode (S3590-08, Hamamatsu), reverse biased at 61.425 V (Keysight). The signal from the photodiode was bandpass filtered and fed into a lock-in amplifier (HF2LI, Zurich Instruments) for image formation. The power of the Stokes beam was set to 15 mW at the sample, and the power of pump was set to 20 mW, and the images were acquired at a size of 512×512 using a pixel dwell time of 10 μs and a lockin time constant of 3 μs.

For two photon fluorescence imaging the output of the OPO was tuned to 900 nm, and delivered to the sample using the pump excitation path described above. The power was set to 20 mW, and a pixel dwell time of 2 μs was used to collect time series from a single point in the center of the image.

### Magnetic bead control in the thin-chip

For the single bead immobilization experiment, 3 μm FlashRed fluorescent dye coated beads (Bangs Labs) were diluted to 0.1% w/v in 1X PBS buffer with 0.5% Tween20, and injected into the thin-chip mounted on the inverted microscope. While imaging the fluorescent beads with 60x water immersion objective and two-photon excitation, the flow pressure was varied from 0.5 to 4.5 PSI. The magnetic field strength was also controlled by moving a stack of ten 10 mm diameter and 2 mm thick neodymium disc magnets mounted on the center of the condenser holder (and field of view).

For the reversible magnetic bead column creation experiment, 1X 1 μm SPRI beads (Beckman) in 1X PBS buffer with 0.5% Tween20 were injected into the thin-chip. While imaging the beads in the flow channel with bright-field microscopy under 5 to 10 PSI flow pressure, the magnet stack was placed as close to the flow channel without blocking the illumination. Four different conditions were imaged, magnet on/off and valve on/off, to test if the magnetic bead column can be formed. To mimic the traditionally fabricated thick chip, we placed a clean 1 cm PDMS block on top of the mounted thin-chip and repeated the experiment.

### Cell culturing

Murine neuroblastoma-derived N2a cells (UC Berkeley cell culture facility) were cultured in Dulbecco’s Modified Eagle Medium (DMEM) with L-glutamine and without pyruvate, including 10% fetal bovine serum (FBS), and 1% penicillin/streptomycin. Cells were passaged upon achieving between 70-100% confluency. Cells were imaged at 50-70% confluency. Cells were transferred to PBS prior to imaging.

