## supplementary materials for "Paper-Thin Multilayer Microfluidic Devices with Integrated Valves"

Supplementary Information

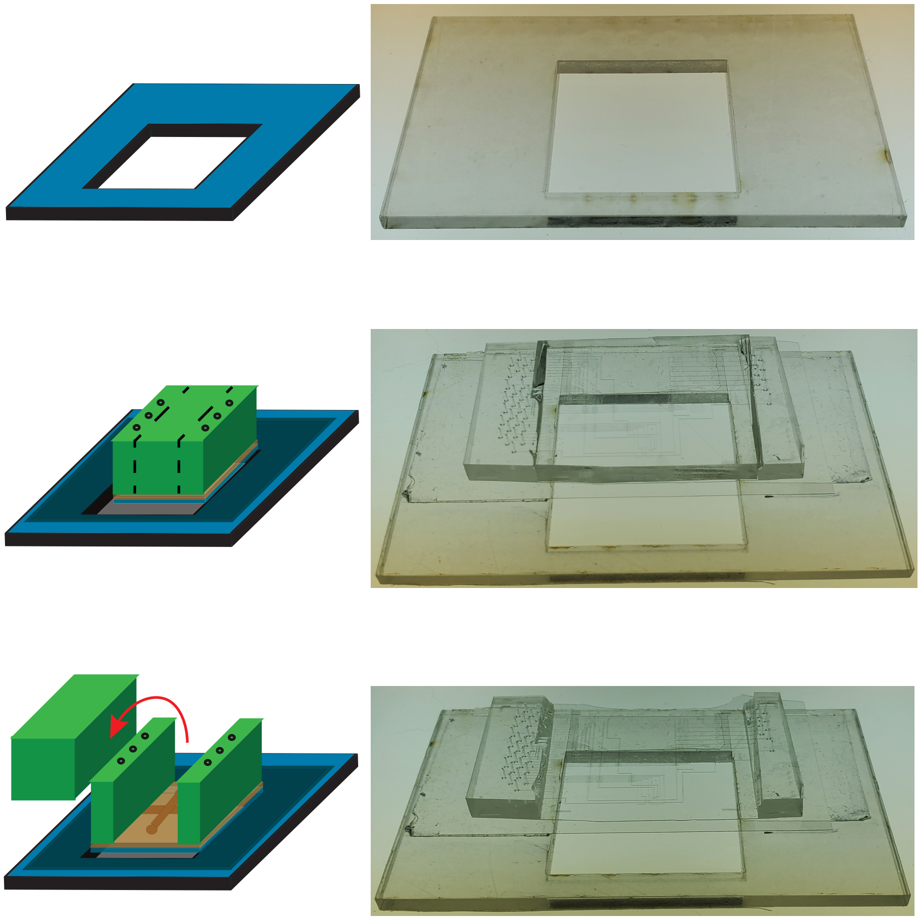

**Supplementary figure 1.** Schematic (left) and photo (right) of the thin-chip mounted on the adapter. The adapter (top) is mounted with the thin-chip on a number 1.5 cover-glass (middle), where the middle part of the carrier is peeled away to expose the thin imaging region (bottom).

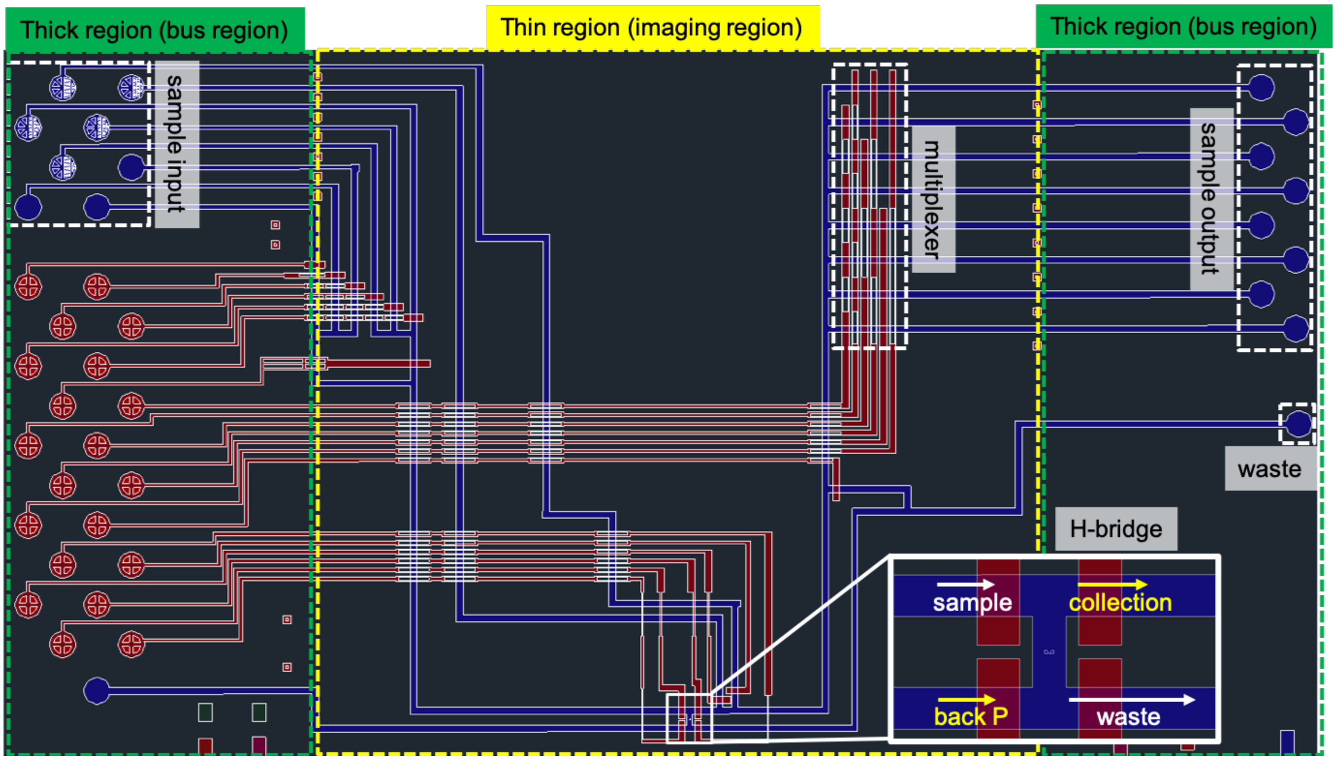


**Supplementary figure 2.** The CAD design of the thin single-cell sorting chip. The chip has sample input and outputs, H-bridge cell imaging and sorter, and a multiplexer (white dotted boxes). The middle area is the imaging region (yellow dotted box) flanked both sides by the thick bus region (green dotted box).

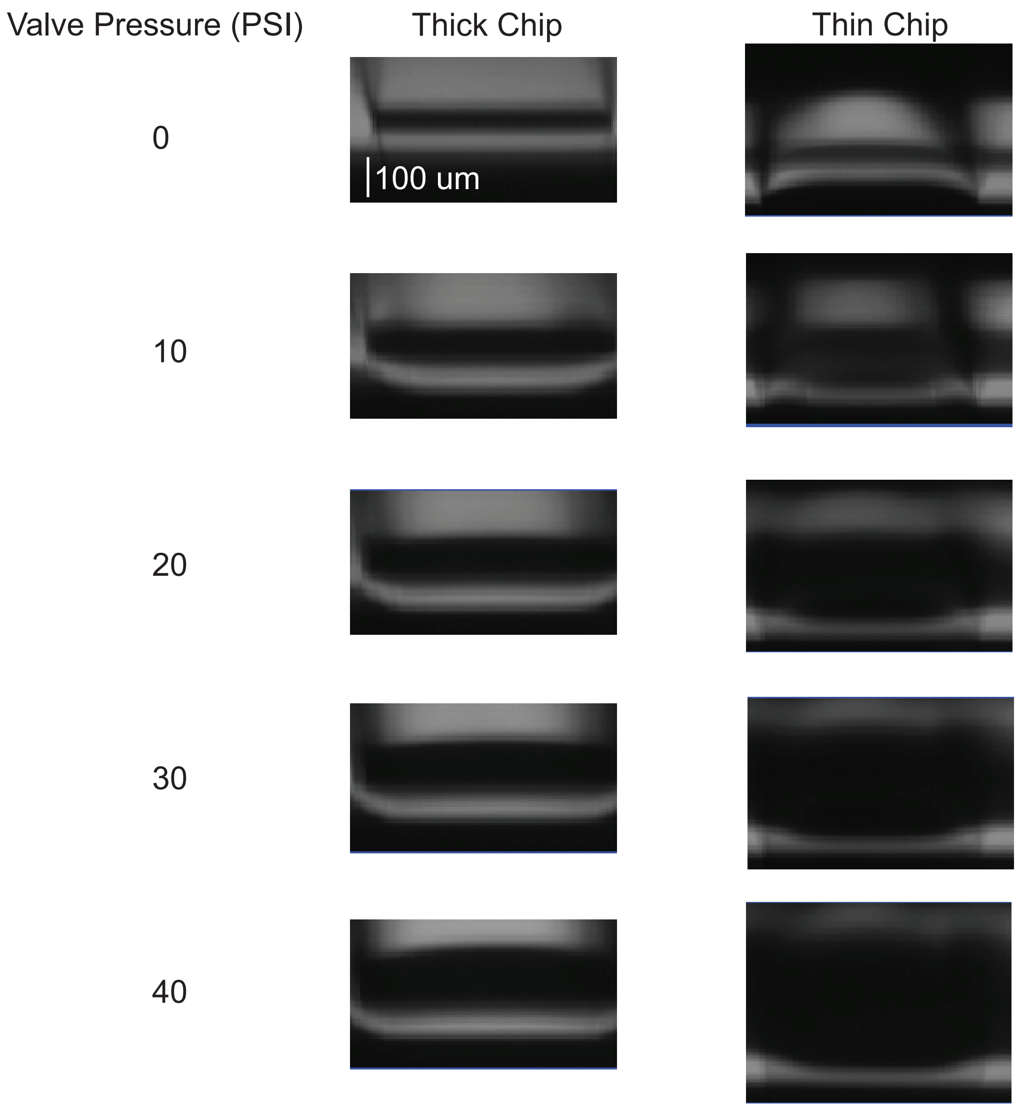


**Supplementary figure 3.** Micro-valve deformation comparison between the thin and thick chip. The vertical cross-section of the micro-valve is obtained by scanning the SRS microscope in xz while gradually increasing the valving pressure.

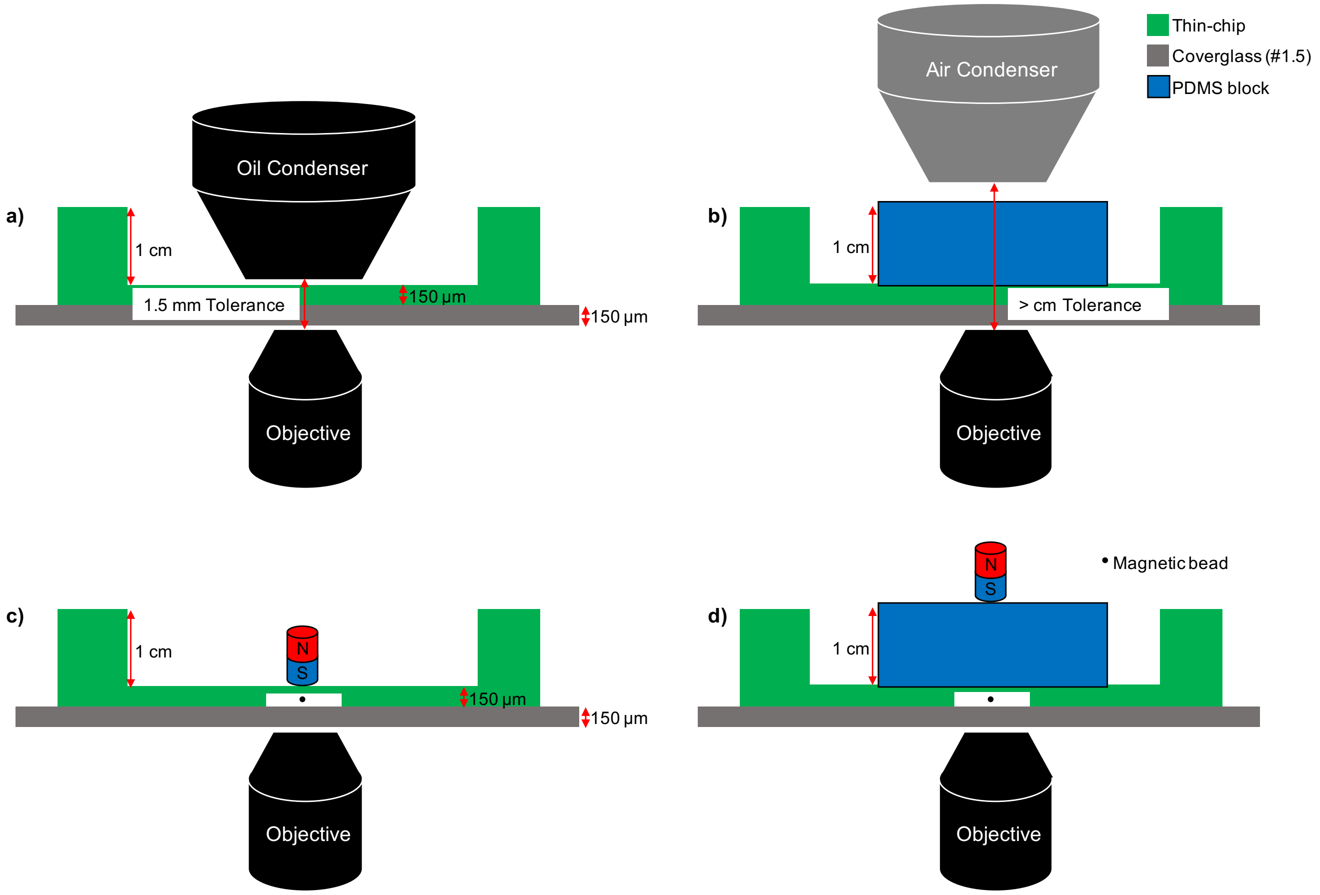


**Supplementary figure 4.** The schematic of the experimental configuration for the thin-chip and thick chip mimic on the inverted microscope. The configuration of SRS cell imaging the **a)** thin-chip with an oil condenser and **b)** the thick chip mimic with an air condenser. The experimental setup for the magnetic bead manipulation experiment on **c)** the thin-chip and **d)** the thick chip mimic. The components, size, and distances are labeled (not to scale).

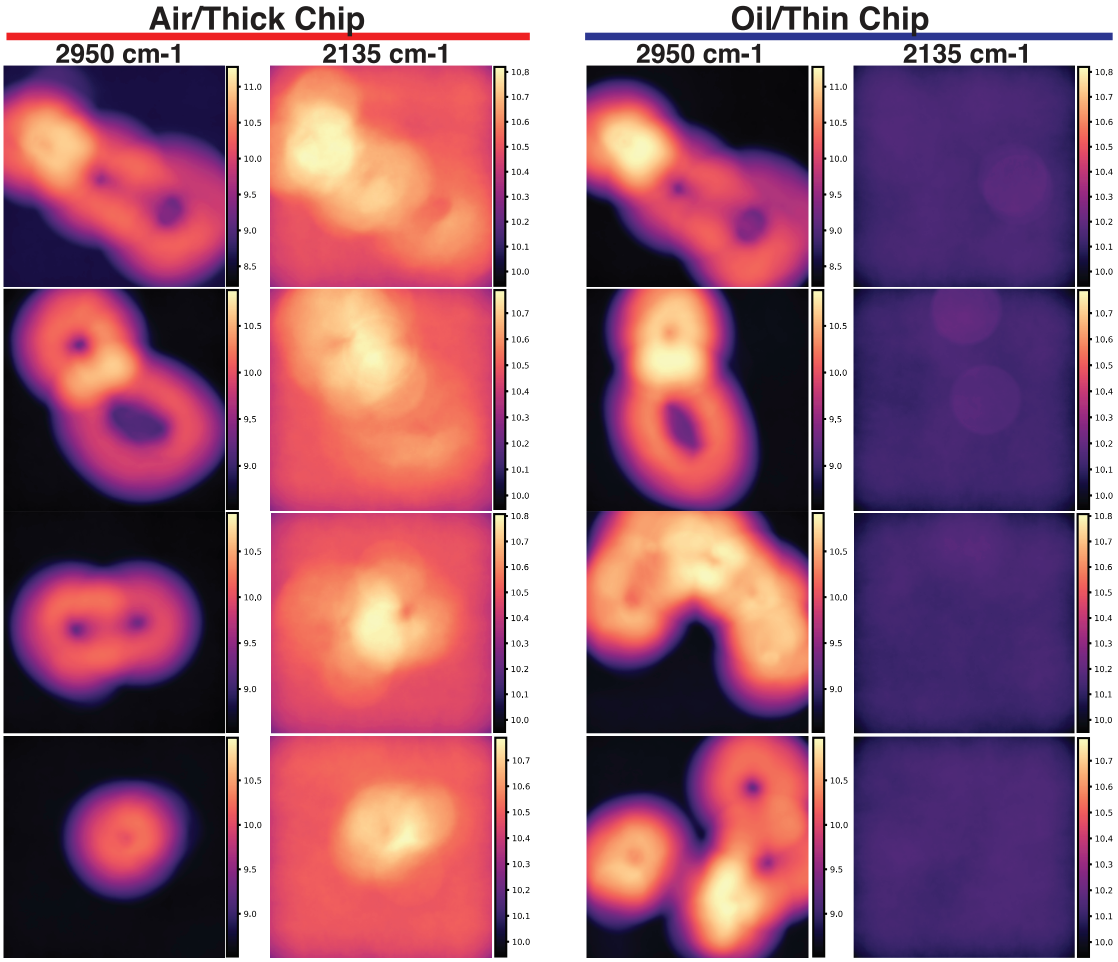


**Supplementary figure 5.** Local entropy maps for mouse L2a cells in the thin and thick chip. The left two columns are pixel entropies from images of cells in a thick chip using a low-NA long working distance air condenser and the right two columns are obtained in the thin chip using a high-NA short working distance oil condenser. Both 2135 $cm^{-1}$ and 2950 $cm^{-1}$ wavelengths acquired.
